# Bend-Induced Twist Waves and the Structure of Nucleosomal DNA

**DOI:** 10.1101/256818

**Authors:** Enrico Skoruppa, Stefanos K. Nomidis, John F. Marko, Enrico Carlon

## Abstract

Recent work indicates that twist-bend coupling plays an important role in DNA micromechanics. Here we show that this coupling induces standing twist waves in bent DNA, and we provide an analytical solution of the minimum-energy shape of a bent double-helical molecule. This solution is in excellent agreement with both coarse-grained simulations of DNA minicircles and experimental structural data for nucleosomal DNA, which is bent and wrapped around histone proteins in a superhelical conformation. Our analysis shows that the observed twist oscillation in nucleosomal DNA, so far attributed to the interaction with the histone proteins, is an intrinsic feature of free bent DNA, and should be observable in other protein-DNA complexes.

## Introduction

Elastic models of DNA have been a key tool for understanding the response of the double helix to applied stresses [1]. Such stresses are ubiquitous in cells, where DNA is continuously being bent and twisted. For instance, in eukaryotes about 75% of the DNA is wrapped around cylindrically-shaped octamers of histone proteins [2]. The 147 base pairs (bp) of wrapped DNA sequence and the histone form the nucleosome, which represents the lowest level of chromosomal organization.

At length scales of a few nanometers the behavior of DNA can be modeled by a homogeneous elastic rod, with stiffness constants associated with the different types of mechanical deformations [3–6]. The simplest such model is the twistable wormlike chain (TWLC), which treats bending and twist as independent deformations. However, symmetry analysis of the right-handed, oppositely-directed-backbone double helix indicates that there must be a coupling of bending to twisting [7]. This can be understood as a consequence of the asymmetry between the major and minor grooves of the double helix. Only a few prior works have considered twist-bend coupling [8–12], and its effect on equilibrium and dynamics of DNA remain largely unexplored.

Here, we show that in bent DNA the twist-bend coupling induces a standing twist-wave at a spatial frequency corresponding to the intrinsic twist of the double helix, *ω*_0_ = 2*π*/(3.6 nm) = 1.75 nm^−1^. This follows from a simple analytical energy minimization of the elastic rod including twist-bend coupling, and from coarse-grained molecular dynamics simulations of a more realistic model of DNA [13]. We also analyze X-ray crystallographic structures of nucleosomal DNA (DNA wrapped around histones), and show that they display twist waves quantitatively matching the predictions of our simple theory. On top of providing a convincing proof of the role played by twist-bend coupling in a key protein-DNA complex (the nucleosome), the generality of our prediction of bend-induced twist waves indicates that similar results should be observable for other protein-DNA complexes.

## Theory and Energy Minimization

Following prior work [7], we describe the double helix centerline using a space curve in arc-length parameterization, with coordinate *s* running from 0 to the total DNA length *L*; we thus treat the double helix as inextensible, which turns out to be appropriate for our purposes. Along the curve we define an orthonormal triad 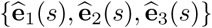, where 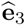 is tangent to the curve, while 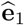 and 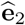 lie on the plane of the ideal, planar Watson-Crick base pairs [7], with 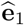 directed along the symmetry axis of the two grooves, pointing in the direction of the major groove. Orthogonality then determines 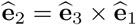 (see Fig. 1).

**FIG. 1.**
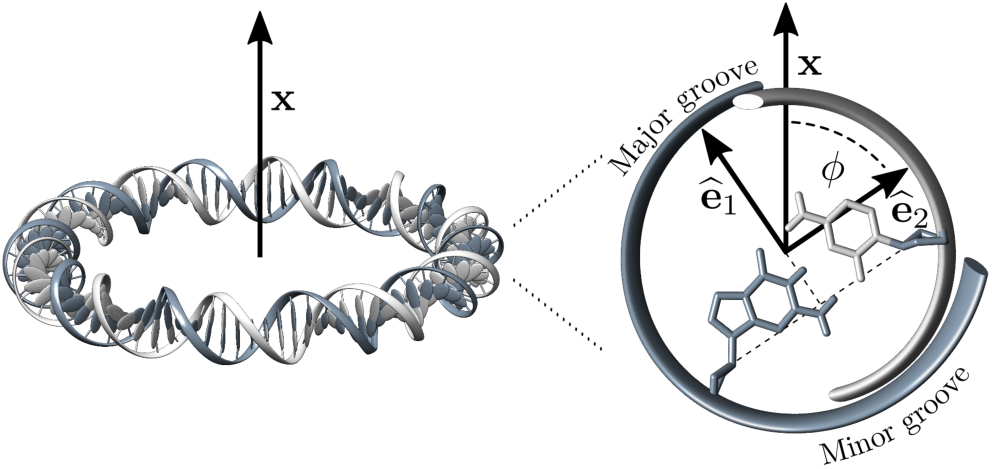
Left: Schematic view of a DNA minicircle lying on a plane orthogonal to a vector x. Right: Zoom-in of a cross-section of the double helix showing the unit vectors 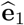 and 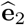 (the tangent vector 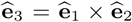 points inside of the page). In an ideal fully-planar circle x lies on the plane spanned by 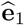 and 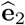. *ϕ* is the angle between 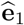 and x.

The three-dimensional shape of the space curve is fully described by the 3-vector field **Ω** that rotates the local unit vectors,

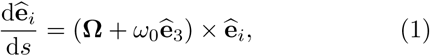

where the index *i* runs over the three spatial directions, and where *ω*_0_ is the intrinsic twist-density of the double helix. As is familiar from mechanics, the rotation vec-tor 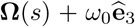 relates the triad at *s* + d*s* to that at *s*. The three components of **Ω**(*s*) along the triad axis are 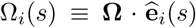. Ω_1_ and Ω_2_ are bending densities (corresponding to the “tilt” and “roll” deformations, respectively, of the DNA literature), with the usual curvature of the backbone given by 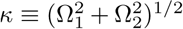. Ω_3_ is the twist density, or, more precisely, the “excess” twist over that of the double helix ground state, *ω*_0_.

Assuming the ground state to be a straight configuration with constant twist density *ω*_0_, one can interpret **Ω** as a strain-field associated with a free energy density. Taking the symmetries of the double helix into account, the deformation free energy to second order in **Ω** is [7]

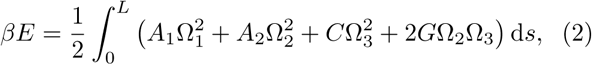

where *β* = 1/*k*_*B*_*T* is the inverse temperature, and *A*_1_, *A*_2_, *C* and *G* are the stiffness parameters. Equation (2) is characterized by a twist-bend coupling term connecting a bending deformation towards the DNA groove (Ω_2_) to a twist deformation (Ω_3_). *G* denotes the twist-bend coupling constant, without which (*G* = 0) one recovers the TWLC.

We investigate the lowest-energy configuration of a circularly-bent DNA molecule, a constraint which can be mathematically imposed by appropriate Lagrange multipliers. This is usually performed by parametrizing Ω_*i*_ in a lab frame using Euler angles (see e.g. Refs. [14, 15]), and numerically solving the corresponding Euler-Lagrange equations. We will instead introduce an approximation, which will allow us to work in the material frame using the Ω’s as minimization variables, and perform the minimization analytically.

One might be tempted to fix the curvature 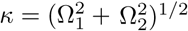 using a Lagrange multiplier, but this leads to a helical solution, rather than a closed configuration [16]. This is a consequence of the bending anisotropy (*A*_1_ ≠ *A*_2_), together with the fact that the plane on which the bending takes place is not restricted. Instead, we seek to impose bending on a plane, as e.g. illustrated in Fig. 1 (left). The bending component of a local deformation is described by the vector 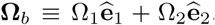. Enforcing bending along a fixed plane, as for instance the plane orthogonal to a vector 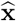, is equivalent to requiring **Ω**_*b*_ to be parallel to 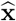. The term 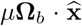 provides a suitable constraint, with *μ* as the Lagrange multiplier. This can be rewritten in the following form

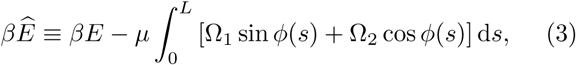

where we have assumed that 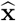 lies on the plane spanned by 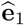 and 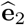, and that *ϕ* is the angle formed between 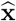 and 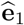 (see Fig. 1). For a straight unbent DNA lying on the 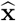 plane (*μ* = 0) we have *ϕ*(*s*) = *ω*_0_*s*. If within one helical turn bending is relatively weak (i.e. *κ* ≪ *ω*_0_), we can approximate *ϕ*(*s*) ≈ *ω*_0_*s*, with the energy minimization then leading to the simple result

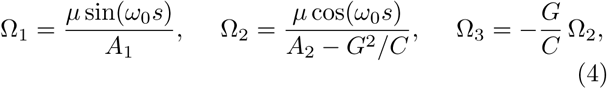

with *μ* ≡ *l*_*b*_/*R*, where *R* is the average radius of curvature and *l*_*b*_ the bending persistence length of the model (2) [11]. The details are discussed in the Supplemental Material [16].

This solution describes a curve with small off-planar periodic fluctuations appearing in the form of standing waves in bending and twist. A non-vanishing *G* is essential for the emergence of twist waves. Although our minimization is not exact, as it is performed under a fixed “background” *ϕ*(*s*), simulations of DNA minicircles of radii ≈ 5 nm (see below, [16]) are in excellent agreement with Eq. (4). In an alternative approach [16] one can obtain twist-waves using a systematic perturbation scheme in powers of *κ*/*ω*_0_, similar to that of Ref. [7]; this parameter is *κ*/*ω*_0_ ≈ (1/5)/1.75 ≈ 0.11 for a DNA mini-circle of radius 5 nm, justifying our approximation [16].

## Coarse-grained DNA simulations

We have performed computer simulations of minicircles with oxDNA, a coarse-grained DNA model in which the double helix is composed of two intertwined strings of rigid nucleotides, held together by non-covalent interactions [13, 17]. Base-pairing together with all other interactions are homogeneous, i.e. sequence-dependent effects are neglected. Various aspects of the mechanics of DNA minicircles, such as kinking, melting and supercoiling, have been discussed in the literature using oxDNA, other coarse-grained models or all-atom simulations [15, 18-21]. Here we focus on the ground-state shape of homogeneous minicircles, and in particular on circular molecules of 85 base pairs (bp), or about 29 nm in length (see Fig. 1). With this choice of length the two ends of the molecule can be joined together without introducing an excess linking number. Two versions of oxDNA were used, see Fig. 2(a,b). In the first version (oxDNA1) the helical grooves have equal width [13], while in the second version (oxDNA2) the grooves are asymmetric, as in real DNA [17]. More details on simulations can be found in Supplemental Material [16].

**FIG. 2.**
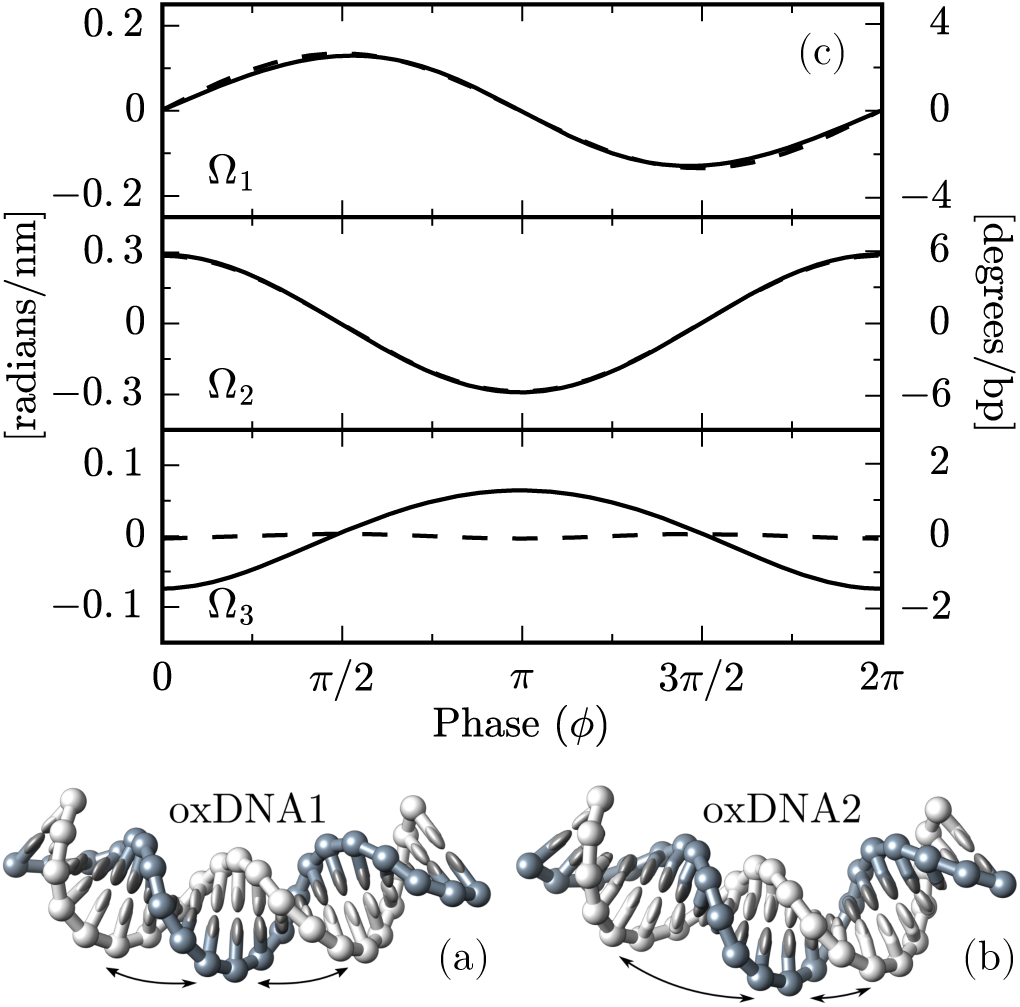
(a,b) Snapshots of minicircles fragments from simulations of oxDNA1 (with symmetric grooves, (a)) and of oxDNA2 (with asymmetric grooves, (b)). (c) Plot of average values of Ω_*i*_ vs. *ϕ* from oxDNA1 (dashed lines) and oxDNA2 (solid lines) simulations. oxDNA2, but not oxDNA1, has a pronounced twist wave. Overall the data are in good agreement with Eqs. (4). A zoom-in of the Ω_3_ for oxDNA1 shows a very weak wave with frequency 2*ω*_0_. This is due to anisotropic bending, as discussed in the Supplemental Material [16]. The Ω_*i*_, as defined in (2), have units of inverse length, which are shown in the left vertical axis. The right axis is in degrees per base pairs, and is obtained by multiplying the left scale by *πa*/180, with *a* = 0.34 nm the base pair separation.

Figure 2(c) shows a comparison between oxDNA1 and oxDNA2 simulations (dashed and solid lines, respectively), in which the Ω_*i*_ are plotted as a function of the base-pair phase angle *ϕ*. The latter was obtained from a Fourier analysis of simulation data: a discrete Fourier transform provides a dominant frequency *ω*_0_ and a global phase *ψ*. From these the local phase of each individual base pair was obtained as *ϕ*_n_ = mod(*ψ* +*naω*_0_, 2*π*), with the index *n* = 0,1… 84 labeling the base pairs along the circle, and *a* = 0.34 nm being the base pair separation. The smooth curves of Fig. 2(c) are obtained by binning the data in *ϕ* and averaging Ω_*i*_ within each bin. A key result of Fig. 2(c) is the clear difference in the behavior of Ω_3_ between the model with symmetric grooves (oxDNA1, dashed lines) and that with asymmetric grooves (oxDNA2, solid lines). The emergent twist waves are associated with the twist-bend coupling interaction [*G* ≠ 0 in Eqs. (4)], which arises from the groove asymmetry of DNA [7]. In the unrealistic case of equal major and minor grooves, one expects *G* = 0, as we indeed observe for oxDNA1. In general, the Ω_*i*_ calculated from oxDNA closely follow the predictions of Eqs. (4). For a quantitative comparison see Supplemental Material [16].

## Nucleosomal DNA

We now turn to the analysis of nucleosomal DNA, which is highly bent around histones, forming a superhelix of radius 4.19 nm and pitch 2.59 nm (for a recent review see e.g. Ref. [2]). The length of the wrapped DNA is 147 bp, corresponding to 1.67 super-helical turns. High-resolution structural crystallographic data for DNA wrapped around histone proteins in nucleosomes is available (we note the seminal work of this type in Ref. [23]). Oscillations in tilt (Ω_1_), roll (Ω_2_) and twist (Ω_3_) were found in early analyses of crystallographic data, and were attributed to histone protein-DNA interactions [23]. Since the publication of the first high-resolution nucleosome data [23], many crystal structures have been determined with different wrapping sequences and various DNA or protein modifications (e.g. methylation and phosphorilation). Here we focus on the average shape of nucleosomal DNA, which can be obtained by averaging over different available structures. Nucleosomal DNA forms a superhelix and not a close circle. Nonetheless, Eqs. (4) are expected to approximate well its shape, as the superhelical pitch is small compared to the intrinsic double-helix twist (details in Supplemental Material [16], see also Ref. [9]).

Figure 3 shows a plot of average Ω_*i*_ vs. *ϕ*, extracted from the analysis of 145 crystal structures from the Protein Data Bank (PDB [24]), using the conformational analysis software Curves+ [22]. The phase *ϕ* is calculated from the discrete Fourier analysis, similarly to the oxDNA data of Fig. 2. From the analysis of crystal structures we find that in nucleosomal DNA Ω_2_ and Ω_3_ have a strong oscillatory behavior for all sequences and are in antiphase as predicted by Eqs. (4). The average of Ω_1_ over all crystallographic data results in a structureless, highly-noisy signal (thin lines, top of Fig. 3). However, a subset of data show oscillations in Ω_1_, detectable from a dominant peak in the Fourier spectrum corresponding to a frequency ≈ *ω*_0_. The average of this oscillating subset is a sinusoidal wave, as expected from Eq. (4). The lack of a clear oscillatory signal may be due to sequence-specific effects and low signal-to-noise ratio, masking the expected behavior.

**FIG. 3.**
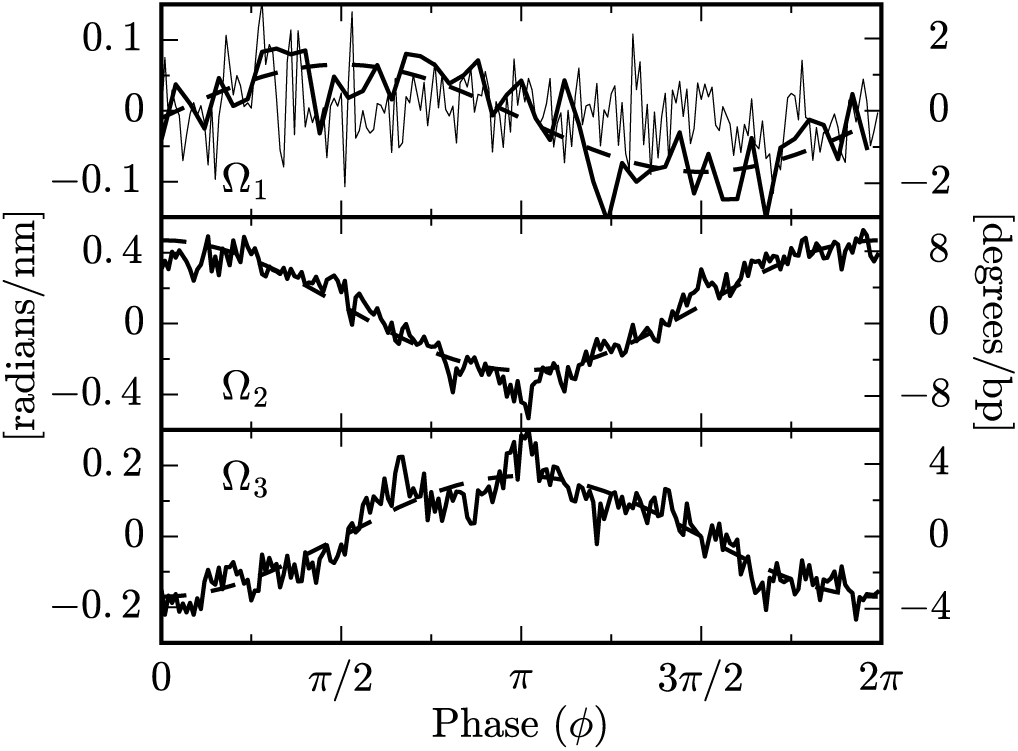
Plot of the mean values of Ω_*i*_ vs. the phase *ϕ* (analogously to Fig. 2), obtained from averaging over 145 nu-cleosome crystal structures. Noisy curves for Ω_2_ and Ω_3_ are simple averages over all structures; smooth curves show the Fourier component for *ω*_0_, indicating its dominance in the average, as well as the antiphase relation of Ω_2_ and Ω_3_ expected from the twist-bend coupling. Data for Ω_1_ averaged over all structures are extremely noisy (light noisy curve), but when selected structures with large power at *ω*_0_ are analyzed (darker curves) the *π*/2-phase-shifted signal expected from theory is observed (see text). The output of the software Curves+ [22] is in degree per bp, given in the right vertical axis.

There is a reasonable quantitative agreement in the wave amplitudes between oxDNA simulations and nucleosome data, as seen by comparing the vertical scales of Fig. 2 and 3. According to Eqs. (4) the wave amplitudes depend on the value of the elastic constants, which may be somewhat different between real DNA and oxDNA. Nucleosomal DNA has a larger amplitude in Ω_2_ and smaller in Ω_1_ than oxDNA. As shown in Supplemental Material [16], from Eqs. (4) it follows that max{Ω_1_} +max{Ω_2_} = 2/*R*, a geometric stiffness-independent constant, *R* being the radius of curvature. Using this relation we find *R* = 4.7 nm both for oxDNA1 and oxDNA2, which agrees with the expected radius *R* = 85*a*/2*π* = 4.6 nm for a 85-bp minicircle. For the nucleosome, we obtain *R* = 4.5 nm, which, considering the large uncertainty on Ω_1_, is reasonably close to the known nucleosomal-DNA radius *R* = 4.2 nm. While the sum of amplitudes Ω_1_ and Ω_2_ is constrained by the geometry, this is not the case for Ω_3_. Its amplitude is larger for the nucleosomal data (Fig. 3) than oxDNA2 (Fig. 2), suggesting that oxDNA2 has a twist-bend coupling constant lower than that of real DNA, in agreement with a previous analysis [12]. From the ratio between the amplitudes of Ω_3_ and Ω_2_ in Fig. 3 and Eq. (4) we estimate *G/C* ≈ 0.46. Recent analysis [11] of single-DNA magnetic tweezers experiments on 7.9 kbp DNA molecules estimated *G* = 40(10) nm and *C* = 110(5) nm, which would yield *G/C* = 0.36(09). Although these two ratios are consistent, some caution is required in their comparison. Simulations have shown that elastic constants for deformations at the base-pair level, relevant for the nucleosome, are generally weaker than stiffnesses for segments of tenths of base-pairs, relevant for the tweezers data [8, 12].

Elastic rod models have been used in the past to investigate various features of nucleosomes [9, 25-29]. In particular, the structure of nucleosomal DNA has been addressed [9] using a model including, besides twist-bend coupling, a stretching modulus and twist-stretch coupling. The elastic energy was minimized while keeping the twist density fixed to the experimentally determined values of Ref. [23], in order to mimic the interaction of DNA with the histone-proteins. In Ref. [29] minimization of a sequence dependent model was performed, while fixing the base pair orientation in 14 known DNA-histones interaction sites [30]. While partially-constraining the conformation of the nucleosomal DNA along the sequence allows for sharper predictions about its local and sequence-dependent behavior, it may obscure some global features. In particular, our work shows that twist oscillations are an intrinsic feature of bent DNA, rather than an explicit consequence of DNA-protein interactions.

## Conclusion

Summarizing, we have shown that in a coarse-grained model of DNA with asymmetric grooves a bending deformation induces an oscillating excess twist having the form of a standing wave. We devised an approximated energy-minimization scheme, which provides analytical predictions for the shape of bending and twist waves. These are in excellent agreement with the numerical simulations, and show that the induced twist waves have a spatial frequency *ω*_0_, the intrinsic DNA twist-density, and an amplitude which is governed by the radius of curvature and the DNA elastic constants. We also showed that crystallographic X-ray nucleosomal DNA data match our prediction of bend-induced twist waves. In nucleosomes, oscillations in DNA twist and bending are usually attributed to the DNA-protein interactions [23], but our work shows that twist waves are general features of bent DNA. We expect that the same kind of correlation will be observed in other protein-DNA complexes, since twist-bend coupling is a fundamental physical property of the double helix.

ES acknowledges financial support from KU Leuven Grant No. IDO/12/08, and SN from the Research Funds Flanders (FWO Vlaanderen) grant VITO-FWO 11.59.71.7N. JM is grateful to the Francqui Foundation (Belgium) for financial support, and to the US NIH through Grants R01-GM105847, U54-CA193419 and U54-DK107980.

